# Biologically Interpretable VAE with Supervision for Transcriptomics Data Under Ordinal Perturbations

**DOI:** 10.1101/2024.03.28.587231

**Authors:** Seyednami Niyakan, Byung-Jun Yoon, Xiaoning Qian, Xihaier Luo

## Abstract

Latent variable models such as the Variational Auto-Encoders (VAEs) have shown impressive performance for inferring expression patterns for cell subtyping and biomarker identification from transcriptomics data. However, the limited interpretability of their latent variables obscures deriving meaningful biological understanding of cellular responses to different external and internal perturbations. We here propose a novel deep learning framework, EXPORT (**EXP**lainable VAE for **OR**dinally perturbed **T**ranscriptomics data), for analyzing ordinally perturbed transcriptomics data that can incorporate any biological pathway knowledge in the VAE latent space. With the corresponding pathway-informed decoder, the learned latent expression patterns can be explained as pathway-level responses to perturbations, offering direct interpretability with biological understanding. More importantly, we explicitly model the ordinal nature of many real-world perturbations into the EXPORT framework by training an auxiliary ordinal regressor neural network to capture corresponding expression changes in the VAE latent representations, for example under different dosage levels of radiation exposure. By incorporating ordinal constraints during the training of our proposed framework, we further enhance the model interpretability by guiding the VAE latent space to organize perturbation responses in a hierarchical manner. We demonstrate the utility of the inferred guided latent space for downstream tasks, such as identifying key regulatory pathways associated with specific perturbation changes by analyzing transcriptomics datasets on both bulk and single-cell data. Overall, we envision that our proposed approach can unravel unprecedented biological intricacies in cellular responses to various perturbations while bringing an additional layer of interpretability to biology-inspired deep learning models.

## 1 Introduction

Transcriptomics data analysis, the study of gene expression patterns across different biological and/or treatment conditions in complex systems, has undergone a transformative journey propelled by advancements in sequencing technologies evolving from bulk to single-cell sequencing to spatial transcriptomics (Li & Wang, 2021; Niyakan et al., 2023). The generated expression profiles are typically used in the context of cell type discovery and cellular development (Møller & Madsen, 2023; Niyakan et al., 2021). Of particular interest are the changes of expression patterns as responses to ordinal perturbations, such as drug screening and radiation exposure at different dosage levels (Peidli et al., 2024; Luo et al., 2022). The cellular responses to external stimuli are intricately linked to the ordinal dosage of the stimulus, delineate dose-response curves that provide invaluable insights into the dynamics of biological systems (Kana et al., 2023). Exploring and understanding how cellular responses evolve across a spectrum of stimulus doses sheds light on the underlying cellular mechanisms and paves the way for precision medicine approaches tailored to individualized dosedependent responses specifically in the context of disease and drug treatment (Bock et al., 2022; Lotfollahi et al., 2019).

Deep generative models such as VAEs (Kingma & Welling, 2014) have shown their capabilities in unraveling biological insights from large and heterogeneous perturbation-induced gene expression profiles due to their model flexibility while keeping necessary changes to the inference procedure relatively minimal (Kana et al., 2023; Lotfollahi et al., 2023). Despite these successes, VAEs suffer from limited interpretability, making them ‘black boxes’ as the reasoning behind predictions as well as the direct correspondence between input features, learned latent vectors and biological processes is often unknown (Lopez et al., 2023). Numerous approaches have been proposed to address the lack of interpretability in deep generative models. For example, disentanglement-promoting VAEs infer disentangled latent representations where one latent unit represents one generative factor of data variability while being invariant to other generative factors (Chen et al., 2018). However, these methods often compromise the quality of the latent variables for downstream analysis tasks (Kimmel, 2020). In computational biology, a recent alternative approach incorporates pathway knowledge to directly modify the VAE architecture, where the neuron connections mirror the user-provided gene-pathway maps and the hidden layers consist of nodes capturing the biological pathway-level activities such as VEGA (Seninge et al., 2021), pmVAE (Gut et al., 2021) and scETM (Zhao et al., 2021) models. (Details in Appendix A.)

Despite the enhanced model interpretability achieved by the pathway informed VAEs when analyzing perturbation-induced transcriptomics data, existing state-of-the-art (SOTA) tools oversimplify the graded biological response dynamics in ordinal perturbations by categorical modeling of stimulus induction levels and ignoring the ordinality (Seninge et al., 2021; Gut et al., 2021). As the perturbation dosage levels are ordered and there are different inter-class importance for each pair of dosage levels, without accounting for the ordered nature of dose-dependent effects, trained models may lead to imprecise prediction of cellular response patterns to ordinal perturbations.

To address current limitations, we propose EXPORT (**EXP**lainable VAE for **OR**dinally perturbed **T**ranscriptomics data), an interpretable VAE model with a biological pathway informed architecture, to analyze ordinally perturbed transcriptomics data. Specifically, the low-dimensional latent representations in EXPORT are ordinally-guided by training an auxiliary deep ordinal regressor network and explicitly modeling the ordinality in the training loss function with an additional ordinal-based cumulative link loss term (Pedregosa et al., 2017). To highlight the capability of our EXPORT in properly encoding perturbed transcriptomics data in an explainable latent space and accurately identifying pathway modules that are differentially affected by dose-dependent perturbations, we have applied EXPORT to two real-world perturbed transcriptomics data by bulk and single-cell sequencing from radiation exposure and chemical induction experiments. Our experimental results demonstrate that EXPORT indeed helps unravel the biological intricacies of cellular response mechanisms to ordinal perturbations.

## 2 Methodology

### 2.1 Model Overview

EXPORT analyzes transcriptomics data (*Y*), under perturbations of the ordinal dosage vector (*D*). It also integrate biological pathway annotations as inputs as indicated in Figures 1.A and B.

**Figure 1:**
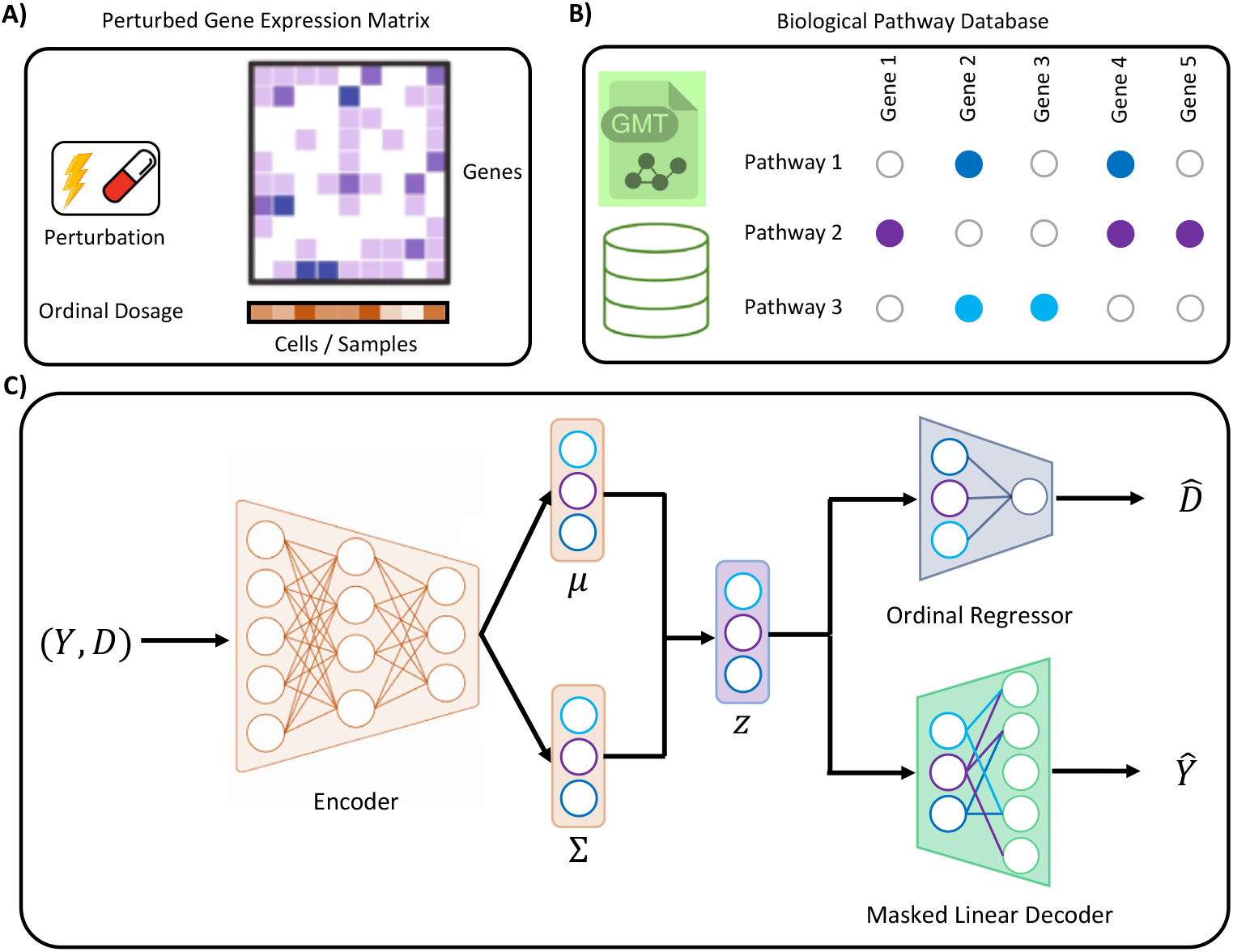
The EXPORT workflow to analyze ordinally perturbed transcriptomics data. EXPORT requires: A) perturbed gene expression data and the ordinal dosage levels of the applied perturbations, as well as B) biological pathway annotations. C) EXPORT model architecture mainly constitutes of a fully connected encoder, a masked linear decoder integrating pathway annotations, and an ordinal regressor network.

#### VEGA Architecture

The encoder-decoder neural network architecture is inspired by VEGA to achieve interpretability leveraging existing gene pathway databases (Seninge et al., 2021). Specifically, the decoder is a sparse single-layer neural network whose wirings are mirroring the user-provided gene-pathway annotation maps to help reconstruct normalized gene expressions, *ŷ*, based on the pathway-constrained latent representations. The encoder is composed of two layers of fully connected nodes with the input number of features being the same as the number of genes in the transcriptomics dataset and the number of its first layer output features set to be 800. The latent space, *Z*, with its dimension set to be the number of extracted pathways from the pathway database, plus one additional fully connected node to capture additional data variability. This choice of the VAE architecture in EXPORT enforces the encoding of the prior biological knowledge that genes work together in coordination in pathways while the deep neural network encoder and decoder capture nonlinear high-order gene-gene interactions (Details in Appendix E).

#### Compositional Decoder

To guide the VAE latent space to capture the intricacies of transcriptomics responses to ordinal perturbations, we integrate an ordinal regessor network taking the lowdimensional latent representations *Z* as inputs to recover the dosage values 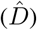. This innovative superposition architecture employs a compositional decoder capable of flexibly incorporating auxiliary information related to the transcriptomics data. In our case, the architecture comprises two decoders: the first decoder *Ŷ* = *D*_1_(*Z*) is dedicated to reconstructing the transcriptomics data, while the second decoder 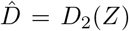 aims at learning the ordinal dosage vector using the same latent representation. Given the independence of each decoder, any available supplementary information can be seamlessly assimilated into the modeling framework.

### 2.2 Training

Following the standard VAE implementations (Kingma & Welling, 2014), the objective to be maximized during training is the evidence of lower bound (ELBO), which is based on the reconstruction loss as well as the Kullblack-Leibler (KL) divergence loss:

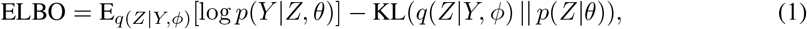

where *ϕ* and *θ* are the learnable variational inference and neural network parameters. Additionally, in EXPORT, as we expect that the latent space *Z* represents a biological pathway activity in the data to analyze, the variational distribution *q*(*Z*|*Y, ϕ*) is modeled as a multivariate normal distribution:

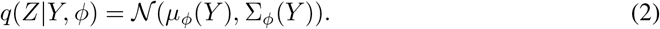

This choice of the variational distribution is common and has proven to work well in previous bulk and single-cell transcriptomics studies (Seninge et al., 2021; Lotfollahi et al., 2019).

#### 2.2.1 Cumulative logistic link model for ordinal regression

In order to explicitly model the ordinal nature of perturbations in real-world experiments to understand transcriptomics responses by guiding the encoding into meaningful pathway-constrained latent representations in EXPORT, we adopt a proportional odds model, one of the first models designed explicitly for ordinal regression problems but lately is recognized as a member of a wider family of models known as cumulative link (CL) models (McCullagh, 1980; Vargas et al., 2020). CL models predict probabilities of groups of contiguous ordinal categories, taking the ordinal scale into account. When analyzing transcriptomics data under ordinal perturbations in EXPORT, assuming that we have *M* different perturbation levels *𝒟* ∈ *𝒟* = {*𝒟*_1_, *𝒟*_2_, …, *𝒟*_*M*_} where there is natural ordering of *𝒟*_1_ ≺ *𝒟*_2_ ≺ … ≺ *𝒟*_*M*_, cumulative probabilities *P* (*𝒟* ≺ *𝒟*_*m*_|*Z*) are estimated, which can be directly related to the following standard probability terms:

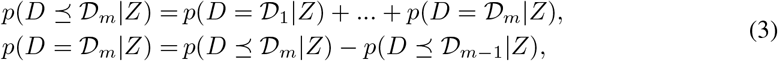

where *m* = 2, …, *M* − 1 and considering that *p*(*𝒟* ⪯ *𝒟*_*M*_ |*Z*) = 1 and *p*(*𝒟* = *𝒟*_1_|*Z*) = *p*(*𝒟* ⪯ *𝒟*_1_|*Z*). The CL model projects the latent variables into a one-dimensional space using the mapping function *f* (·) and then a set of thresholds {*c*_0_, *c*_1_, *c*_2_, …, *c*_*M−*1_, *c*_*M*_}, where *c*_0_ = −∞ and *c*_*M*_ = +∞, estimated from the data, is used to partition the projection into the different ordinal levels. Specifically, the dosage level D_*m*_ is predicted if and only if *f* (*Z*) ∈ [*c*_*m−*1_, *c*_*m*_]. Finally, in the proportional odds model or alternatively the cumulative logistic link model used in our EXPORT framework, we have the following general model form:

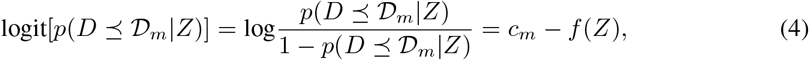

where *m* = 1, …, *M* − 1. Lastly, the cumulative logistic link model loss function is given by its negative likelihood, that is,

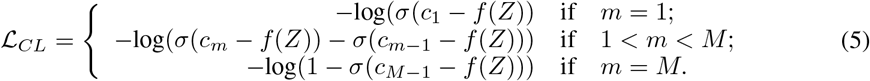

The overall training loss for the pathway-informed supervised VAE model in EXPORT can be derived by integrating the VAE reconstruction loss term in Equation (1) and the ordinal-based cumulative link loss expressed in Equation (5) as the supervised learning to guide the learning of dose-dependent transcriptomics response to perturbations:

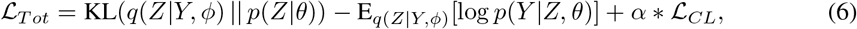

where *α* is a coefficient hyperparameter on the cumulative link loss controlling the effect of ordinal deep regression in training the overall model in EXPORT. The training methodology proposed herein can be conceptualized as an extension of the VEGA model, characterized by its generalization capabilities. Specifically, when the hyperparameter *α* is set to 0, the training paradigm asymptotically aligns with the unsupervised learning framework established by the VEGA model. Conversely, a non-zero value of *α* introduces the capability to allow the ordinal characteristics of perturbations to guide the learning of deep latent representations to better reflect cellular responses to perturbations. This flexibility enhances the model’s ability to capture and represent complex data structures effectively.

## 3. Results

In our experiments with EXPORT model, we apply it to two real-world transcriptomics datsets on both bulk and single-cell resolutions ordinally perturbed with radiation exposure and chemical induction. In the following paragraphs we present our main findings:

### EXPORT latent space captures ordinality of transcriptomic response to radiation exposure

In our first experiment, we investigate the capability of EXPORT in resolving the human cell line response to radiation exposures at different levels. For this purpose, we analyze the publicly available radiation exposure microarray gene expression data (Nosel et al., 2013) (Detailed in Appendix B).

The decoder neural connections in our EXPORT model, mirror the gene-pathway mappings in the KEGG database when analyzing this radiation exposure data (Appendix B). After mapping the actual radiation dosage values of 121 samples in the dataset to the ordinal labels from 0 to 6, we train an EXPORT model for 200 epochs to embed the gene expression data into a ordinality-preserving and interpretable lower-dimensional pathway-constrained latent space. Figures 2.A and .B display the UMAP (Uniform Manifold Approximation and Projection) (McInnes et al., 2020) embedding of the derived latent representations by the corresponding EXPORT model. These visualizations illustrate that EXPORT’s learned latent space effectively captures the gradual changes in cellular transcriptomic response to radiation exposure across different ordinal levels with embedded points ordered hierarchically based on assigned ordinal labels.

**Figure 2:**
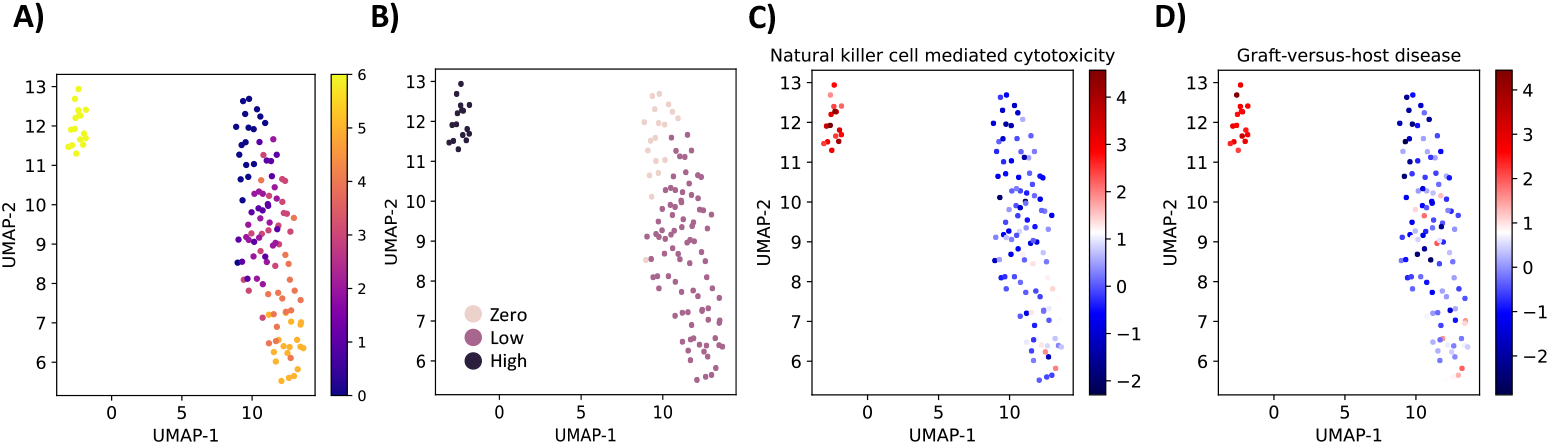
UMAP embedding of the latent representations learned by the EXPORT model trained on the radiation exposure data with the samples colored by A) ordinal dosage labels, B) dosage levels, and C-D) EXPORT-inferred activity heatmaps of two common top differentially activated pathways in the comparison of zero versus low and high dosage level samples.

Furthermore, to identify the KEGG pathways that are differentially activated under either low-dose or high-dose radiation exposure, compared to samples with no radiation exposure, we have applied the Bayesian hypothesis testing procedure that has been implemented in EXPORT as described in Appendix C. Intersecting the top 5 differentially activated pathways in zero versus low-dose and high-dose exposure sample groups respectively, we find two common KEGG pathways: *Natural killer cell mediated cytotoxicity* and *Graft-versus-host disease*. Both pathways have been reported previously as pathways involved in radiation exposure experiments confirming the reliability of EXPORT in inferring the pathway activity scores (Luo et al., 2022). Notably, these two pathways ranked low in the unsupervised version of EXPORT that does not integrate the ordinal dosage labels when deriving latent representations of pathway responses, highlighting the importance of the ordinal regression module in analyzing ordinally perturbed transcriptomics data. (Detailed in Appendix D.1).

### Ablation study for the ordinal regression module in EXPORT

To assess the ordinal regressor network’s effectiveness in capturing ordinality of dose-dependent responses in the EXPORT derived latent space, we conduct detailed ablation experiments, encompassing the modifications of the cumulative link loss hyperparameter coeffcient *α* in Equation (6). As depicted in Figure 3, the results indicate that EXPORT is unable to capture any ordinal structure in the latent space with an *α* = 0, which is equivalent to the unsupervised learning framework established by the VEGA model (Seninge et al., 2021). Furthermore, we observe that gradually increasing the *α* hyperparameter in the EXPORT loss function helps the corresponding trained models to resolve the inherent ordinality in dose-dependent transcriptomics response to radiation exposures, highlighting the necessity of have the ordinal regressor network and the cumulative link loss for ensuring inference of ordinal-preserving latent space by EXPORT. (Additional ablation study on ordinal regression network replacement with a classifier one is presented in Appendix D.2)

**Figure 3:**
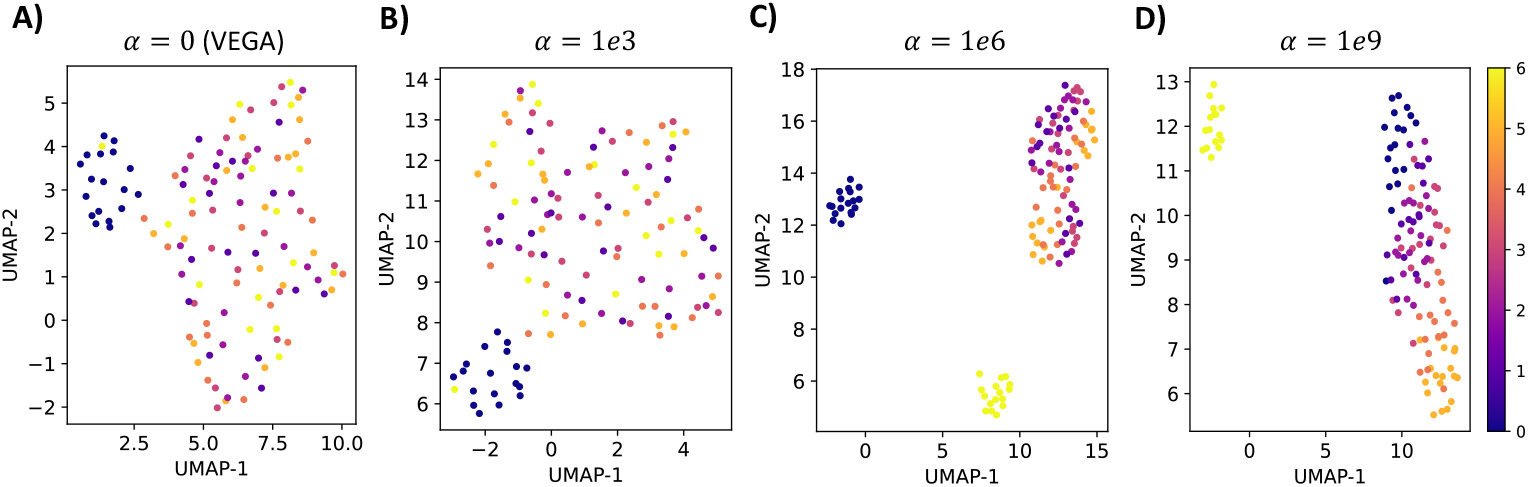
Ablation study on the ordinal regression module in EXPORT.

### EXPORT resolves the hepatic response to ordinal TCDD induction at single-cell resolution

To further assess EXPORT’s ability to capture interpretable and ordinally preserved low-dimensional representations of large single-cell transcriptomics datasets, we apply it to chemically perturbed single-nuclei RNA sequencing (snRNA-seq) data from mouse liver cells (Details in Appendix B) (Nault et al., 2022). The dataset includes mouse liver cells treated with varying doses of 2,3,7,8-tetrachlorodibenzo-p-dioxin (TCDD). Our goal is to understand dose-dependent responses of hepatocyte-portal cells among other cell types in the dataset. After preprocessing and assigning ordinal dosage labels, we train an EXPORT model with decoder neural connections reflecting gene-pathway relationships from the mouse Wikipathways database (Martens et al., 2020) for 50 epochs (Appendix B). Figure 4.A depicts the UMAP embedding of learned latent representations by EXPORT, organized based on ordinal TCDD-induction dosage labels, showcasing EXPORT’s ability to unveil ordinally preserved and biologically meaningful latent spaces in this large-scale perturbed transcriptomics dataset at single-cell resolution.

**Figure 4:**
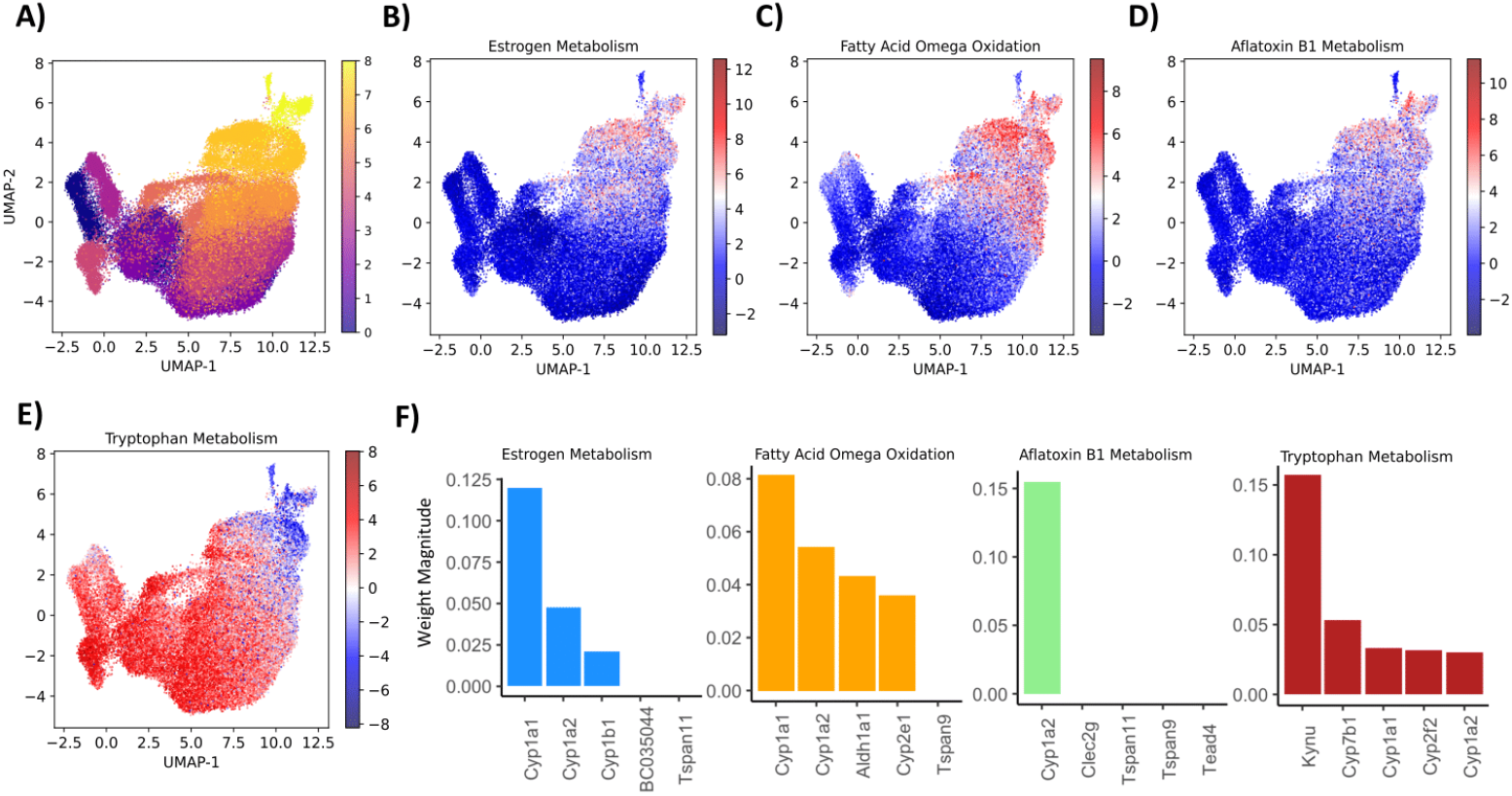
UMAP embedding of the latent representations learned by the EXPORT model trained on the TCDD-induction snRNA-seq data with the cells colored by A) ordinal dosage labels, and B-E) EXPORT-inferred activity heatmaps of four common top differentially activated pathways in the comparison of high versus low and zero dosage level samples. F) Top five genes with the highest decoder weight magnitudes in each of the pathways.

Similar to our radiation exposure data analysis experiment, we group the cells into three different categories of high, low and zero TCDD induction levels (Appendix B) and perform differential pathway activity analysis for the results obtained by EXPORT (Appendix C). Intersecting the top 5 differentially activated pathways in high versus zero-dose and low-dose TCDD induction sample groups respectively, we find four common pathways: *Estrogen metabolism, Fatty acid omega oxidation, Aflatoxin B1 metabolism* and *Tryptophan metabolism*. All these four pathways have been previously reported as known well-established pathways involved in hepatic responses to TCDD in mice such as Cholico et al. (2021), Friedrich et al. (2021), Zhu et al. (2008) and Zhu et al. (2021) supporting evidences for activation of *Fatty acid omega oxidation, Tryptophan metabolism, Estrogen metabolism* and *Aflatoxin B1 metabolism* pathways after TCDD induction.

Additionally, using EXPORT’s interpretability within the decoder structure, we examine the inferred decoder weights to highlight genes’ impact on pathway activity scores. Specifically, we extract the top five genes with the highest weight magnitude for the four common differentially activated pathways. In Figure 4.F, we observe well-known markers of TCDD-induced hepatotoxicity, such as genes from the cytochrome P450 family (Cyp1a1 and Cyp1a2), repeatedly appearing among the top five genes with the highest EXPORT decoder weights (Zhu et al., 2021; Lindros et al., 1997).

In summary, EXPORT effectively captures validated biological insights from inferred model parameters, demonstrating its ability to analyze perturbed single-cell transcriptomics data and capture cellular dose-response patterns.

## 4 Conclusions

This study introduces EXPORT, a novel interpretable and ordinality-preserving VAE model for embedding ordinally perturbed transcriptomics data. Compared to existing pathway-informed VAE models, EXPORT infers a more ordinality-aware latent space through auxiliary ordinal regressor network training. One limitation is its focus on single transcriptomics data modality. Future research will explore EXPORT’s potential for multi-omics data integration to uncover unprecedented cellular dose-dependent response intricacies. Despite this limitation, our extensive experiments demonstrate EXPORT’s capability in capturing biologically significant regulatory mechanisms, making it a robust and interpretable model to accelerate perturbation biology development.

## A Related work

### VEGA (Seninge et al., 2021)

The VAE enhanced by gene annotations (VEGA) model utilizes prior knowledge of gene pathway modules for designing a sparse linear decoder network to obtain an interpretable latent space in VAEs and subsequently detect active pathway modules from models trained on perturbed transcriptomics data. Despite addressing the interpretability concern, this model treats perturbations as categorical variables and overlooks the ordinal nature of perturbations.

### pmVAE (Gut et al., 2021)

Similar as the VEGA model architecture, pmVAE has masked linear decoder networks to achieve interpretability. However, unlike VEGA, pmVAE trains multiple VAEs, each incorporating prior information of one pathway module for detecting changes to stimuli. As in VEGA, pmVAE oversimplifies the estimation of dose-dependent responses by ignoring the ordinal relationships of perturbations.

### scETM (Zhao et al., 2021)

The single-cell embedded topic model (scETM) uses topic modeling to adjust for potential batch effects and utilizes an encoder network that infers cell type mixtures and a linear decoder based on matrix tri-factorization that incorporates pathway knowledge to discover interpretable cellular response embeddings while integrating single-cell transcriptomics data from different treatment conditions. scETM suffer from lack of ordinal adjustments when analyzing ordinally perturbed transcriptomics data and thus, hierarchical ordinal relationships are also lost in the inferred latent space.

## B Dataset details

For benchmarking and validating our proposed EXPORT model, we use two transcriptomics datasets with different characteristics. The first dataset encompasses microarray gene expression data from human cell lines exposed to a varying range of radiation doses. In our second experiment we select a transcriptomics dataset at single-cell resolution from mouse liver cell lines treated with 2,3,7,8tetrachlorodibenzo-p-dioxin (TCDD) at different dosage levels. In this section, we provide characteristic details of the two transcriptomics datasets analyzed with EXPORT as well as the pathway databases chosen in this manuscript:

### B.1 Transcriptomic datasets

#### B.1.1 Radiation exposure dataset

The radiation exposure dataset is derived from a study conducted by Nosel et al. (2013) encompassing microarray gene expression data from human cell lines exposed to varying radiation doses.

This dataset is publicly available in the Gene Expression Omnibus (GEO) database with the accession number GSE43151. GSE43151 consisting of a total of 121 blood samples, where five healthy male donors provided 400 mL venous peripheral blood samples each. These conditions cover a wide range of radiation doses, spanning from low to high intensities. Before we conduct the ordinal dose-response analysis using EXPORT, the dataset is preprocessed by R `GAGE` package (Luo et al., 2009). The preprocessing steps include filtering and normalizing all 121 samples present in the dataset. Specifically, in the filtering step, we filter out probes that were undetected in at least 75% of the samples which leads us to having total 10, 875 probes after the filtering step. These preprocessing steps are crucial for ensuring a robust and reliable dataset for subsequent analyses with EXPORT and enabling more accurate identification of relevant molecular mechanisms governing the radiation dose-response in human cell lines.

This radiation exposure gene expression data encompasses a spectrum of doses spanning from 0.005 Gy to 0.5 Gy. In order to perform differential pathway activity analysis using EXPORT on this dataset, following the instructions described in Luo et al. (2022), we have categorized samples based on their radiation exposure levels to three main sets of ‘zero radiation’, ‘low-dose radiation’ (0.005 Gy to 0.1 Gy), and ‘high-dose radiation’ (0.5 Gy). Table 1 summarize the radiation exposure gene expression data characteristics.

**Table 1:** Summary statistics of the radiation exposure gene expression dataset GSE43151.

| Dose | Dose Level | Ordinal Dose Label | Number of Samples |
| --- | --- | --- | --- |
| 0 Gy | Zero | 0 | 18 |
| 0.005 Gy | Low | 1 | 16 |
| 0.01 Gy | Low | 2 | 18 |
| 0.025 Gy | Low | 3 | 18 |
| 0.05 Gy | Low | 4 | 17 |
| 0.1 Gy | Low | 5 | 18 |
| 0.5 Gy | High | 6 | 16 |

#### B.1.2 TCDD dose-dependent response single-nuclei rnaseq (snrna-seq) dataset

To showcase the capability of EXPORT in unraveling the ordinal intricacies of perturbed transcriptomics data on single-cell resolution, we select the single nuclei RNA sequencing (snRNA-seq) data of mouse liver cells gavaged with 2,3,7,8-tetrachlorodibenzo-p-dioxin (TCDD) at doses of 0.01, 0.03, 0.1, 0.3, 1, 3, 10, or 30 μg/kg (Kana et al., 2023; Nault et al., 2022). This dataset is publicly available in the Gene Expression Omnibus (GEO) under the accession number GSE184506. A total of 131,613 nuclei are sequenced in this dataset. As due to the uneven expression of the TCDD canonical receptor, the aryl hydrocarbon receptor (AhR), the hepatic responses to TCDD vary across different cell types in the liver and also within cell types (such as hepatocytes), in our analysis with EXPORT, we only consider cells from hepatocyte-portals (Kana et al., 2023). This leads to selection of 57,284 nuclei expression profiles from hepatocyte-portal cell types.

Following the instructions in Kana et al. (2023), we preprocess the data using the `scanpy.pp` package. Specifically, the cell expression vectors are normalized to the median total expression counts for each cell. Then, the cell counts are then log transformed with a pseudo-count of 1 and finally, we select the top 5,000 most highly variable genes. Figure 5.A shows the UMAP embedding of all hepatocyte-portal cells after preprocessing steps colored by their corresponding TCDD dose-induction levels. Additionally, the distribution of cell numbers across different TCDD dosage levels is demonstrated in Figure 5.B. Additionally, Table 2 summarizes the TCDD induction gene expression data characteristics.

**Table 2:**
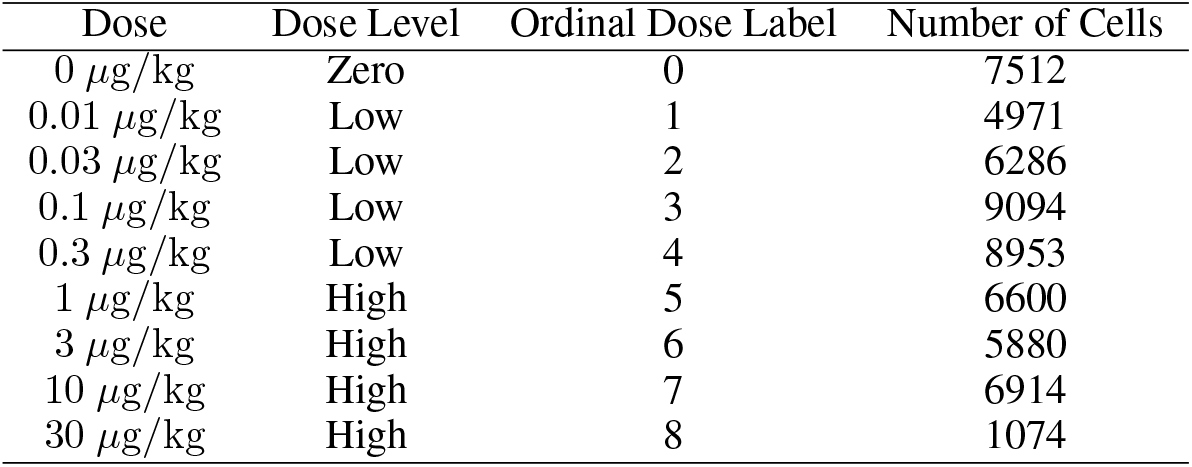
Summary statistics of the hepatocyte-portal cells in TCDD induction snRNA-seq dataset (GSE184506).

**Figure 5:**
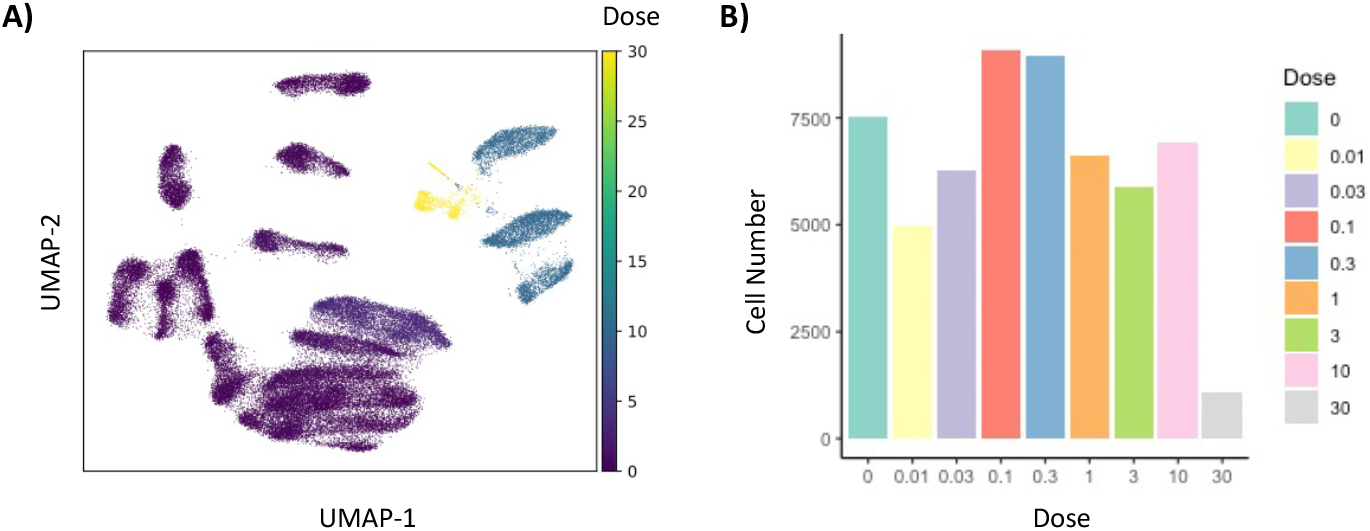
Hepatocyte-portal cells characteristics extracted from the snRNA-seq data perturbed with TCDD at different dosage levels. A) The UMAP embeddings of the preprocessed hepatocyte-portal cells gene expression profiles colored by their corresponding induced TCDD dosage levels. B) Bar plot visualization of the hepatocyte-portal cell number distribution across different dosage levels.

### B.2 Pathway datasets

#### B.2.1 Kegg database for radiation exposure data analysis

Following the instructions in Luo et al. (2022), we have used the KEGG (Kyoto Encyclopedia of Genes and Genomes) database to obtain a reliable set of known biological pathways to enhance our EXPORT model’s explainability when analyzing the radiation exposure gene expression data (Kanehisa & Goto, 2000). KEGG is a collection of manually drawn pathway maps for understanding high-level functions and utilities of biological systems. In our case, we have identified 343 pathways relevant to the gene expression dataset GSE43151 from the available 548 KEGG pathway maps by discarding the pathways that do not contain any gene whose measurement was included in GSE43151 radiation exposure gene expression dataset.

#### B.2.2 Wikipathway database for tcdd-induction data analysis

When analyzing the single-cell snRNA-seq gene expression data of mouse liver cells perturbed with TCDD induction, we have used the 2024 mouse Wikipathways database to form the neural connections in the EXPORT decoder structure (Martens et al., 2020). Specifically, the 2024 mouse Wikipathways database consists of 202 pathways, which we have used their gene-pathway mappings to mask the EXPORT VAE decoder *D*_1_ as described in the methodology section of the manuscript.

## C Differential pathway activity analysis

Differential pathway activity analyses are often of interest when analyzing perturbed transcriptomics datsets of different treatment levels. Inspired by the Bayesian hypothesis testing procedure described in Seninge et al. (2021) and Lopez et al. (2018), for the differential pathway activity analysis, we implement a Bayes factor (BF) (Held & Ott, 2018) based hypothesis testing procedure in EXPORT. Specifically, the posterior probabilities of mutually exclusive hypotheses are approximated through repeated Monte Carlo sampling of the correspondingly derived EXPORT’s latent variable distributions. Then, BF values, the ratio of the hypothesis posteriors, are estimated to rank the pathways differential activities. The sign of the corresponding BF indicates which of the null and alternative hypotheses is more likely, and its magnitude represents the significance level of the pathway differential activity.

## D Additional results

### D.1 Supervised vs. unsupervised modeling

In this section, we provide additional differential pathway activity analysis results associated with EXPORT and its simpler unsupervised version that does not incorporate the ordinal dosage labels when analyzing the radiation exposure data. Figures 6 and 7, demonstrate the top 5 differentially activated pathways when comparing zero versus high and low dosage sample groups respectively. Additionally, EXPORT-derived Bayes factor values are indicated in the figures to show the significance level of the differential activity for the corresponding pathways.

**Figure 6:**
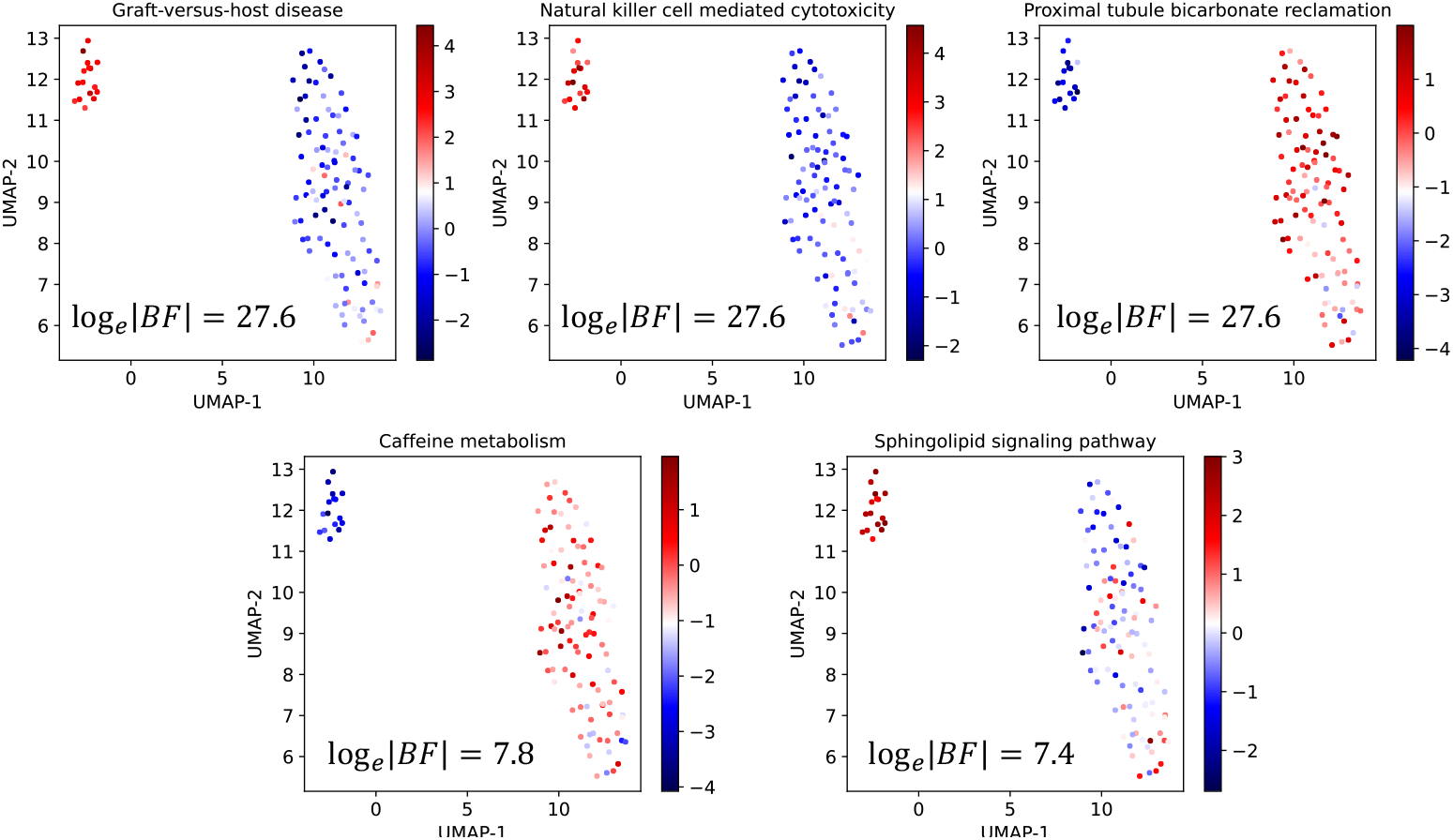
Top 5 identified differentially activated pathways with EXPORT in zero vs high radiation experiments with EXPORT.

**Figure 7:**
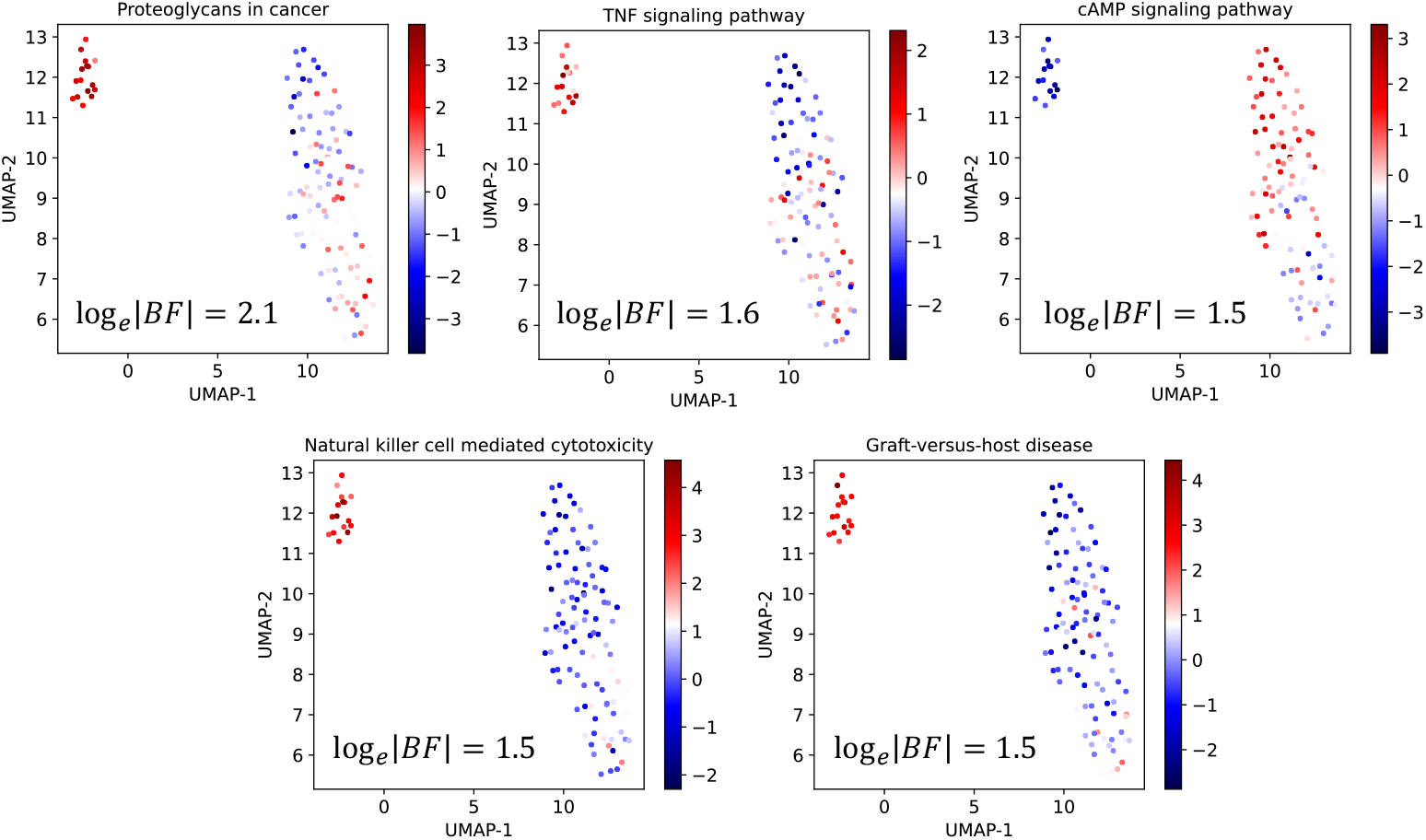
Top 5 identified differentially activated pathways with EXPORT in zero vs low radiation experiments with EXPORT.

Furthermore, to showcase the necessity of ordinal regressor module presence in the EXPORT model when analyzing the ordinally perturbed transcriptomics data, we repeat the differential pathway activity analysis with the unsupervised version of EXPORT that does not incorporate the ordinal dosage labels during the model training. Similarly, after training the unsupervised model we compare zero dosage level group versus low and high dosage level sample groups using the differential activity analysis implemented in EXPORT. Figures 8 and 9, demonstrate the top 5 differentially activated pathways when comparing zero versus high and low dosage sample groups with EXPORT unsupervised version respectively.

**Figure 8:**
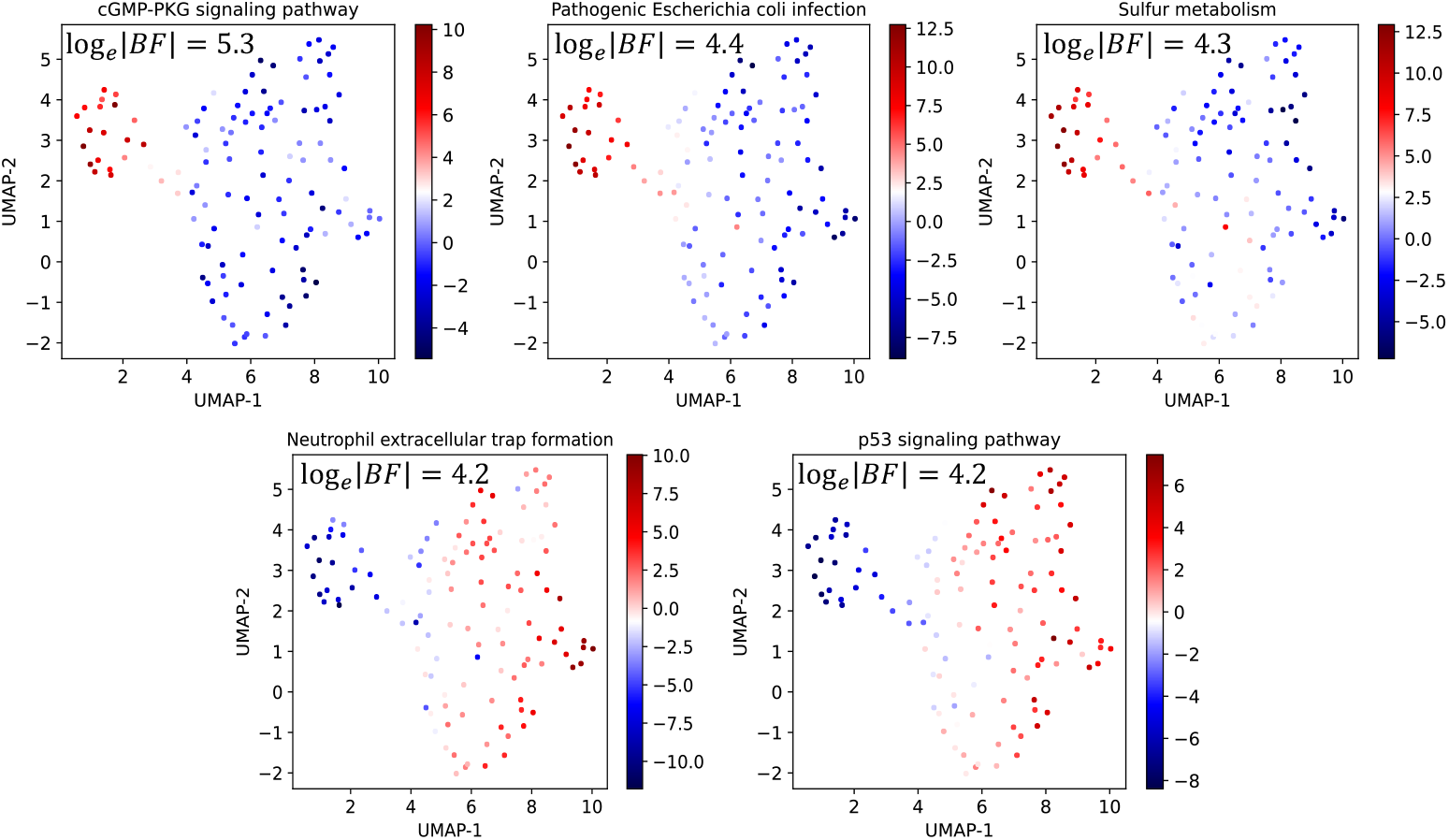
Top 5 identified differentially activated pathways with EXPORT in zero vs high radiation experiments with EXPORT’s unsupervised version.

**Figure 9:**
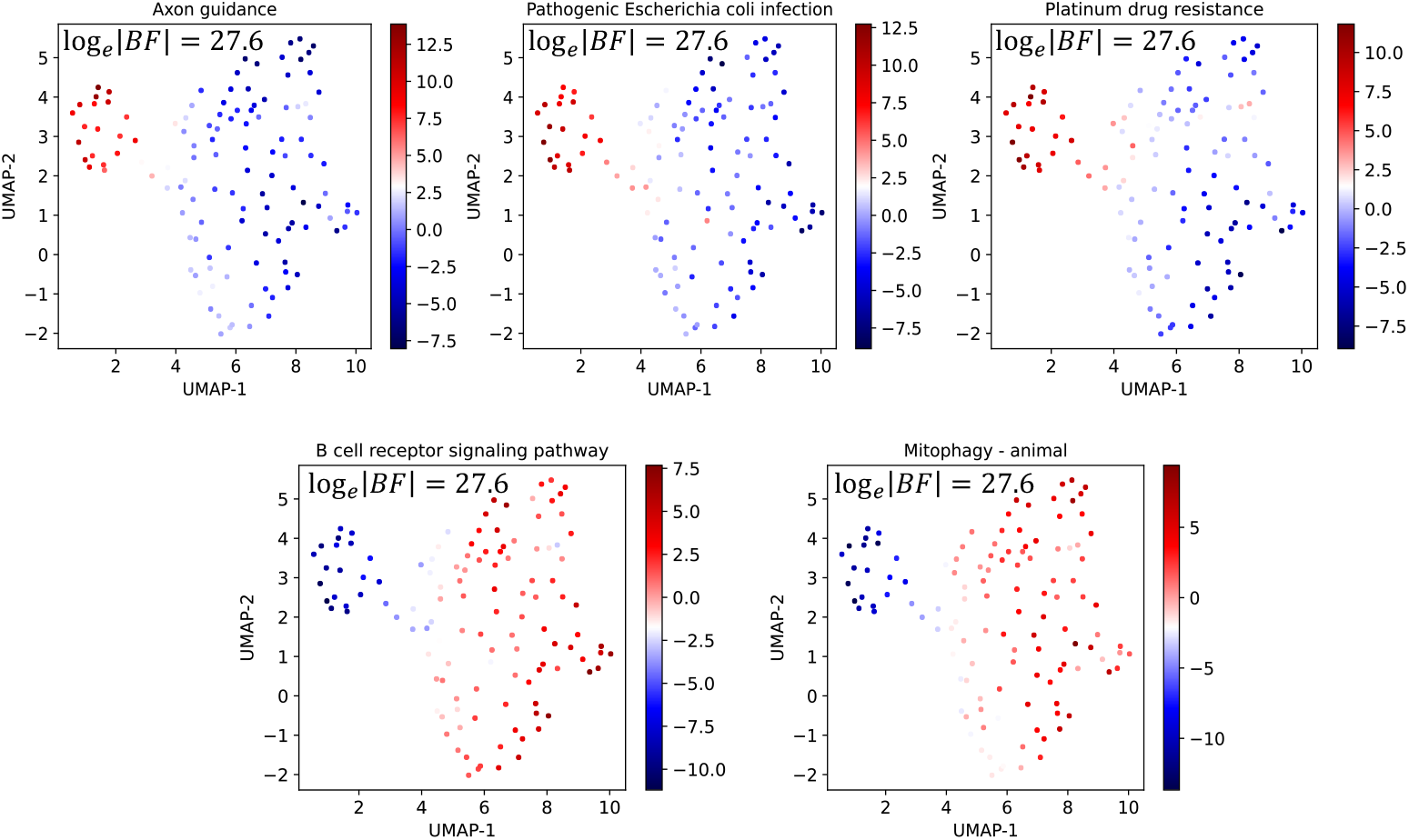
Top 5 identified differentially activated pathways with EXPORT in zero vs low radiation experiments with EXPORT’s unsupervised version.

By investigating the differntial pathway activity results of the unsupervised version of EXPORT, we notice that two KEGG pathways *Natural killer cell mediated cytotoxicity* and *Graft-versus-host disease* which have been previously reported as pathways involved in the radiation exposure experiments (Luo et al., 2022) are ranked low between KEGG pathways. Specifically, the *Natural killer cell mediated cytotoxicity* and *Graft-versus-host disease* pathways are ranked 71 and 209 in zero versus low dosage level sample groups comparison and ranked 149 and 210 in zero versus high dosage level sample groups comparison based on calculated Bayes factor values among all 343 KEGG pathways included in this study highlighting the need for the ordinal regressor module and its corresponding cumulative link loss implemented in EXPORT to achieve reliable results.

### D.2 Ordinal regression vs. multi-class classification

To highlight the importance of the ordinal regression network and the cumulative link loss function utilized in EXPORT, we investigate an EXPORT alternative version in which the ordinal regressor network and ordinal-based cumulative link loss are replaced with classifier network and cross entropy loss function to do multi-class classification task that does not preserve ordinality of perturbation dosage levels. Figure 10 visualizes the overall workflow of this alternative version of EXPORT. For training this deep learning model we utilize the below loss function:

**Figure 10:**
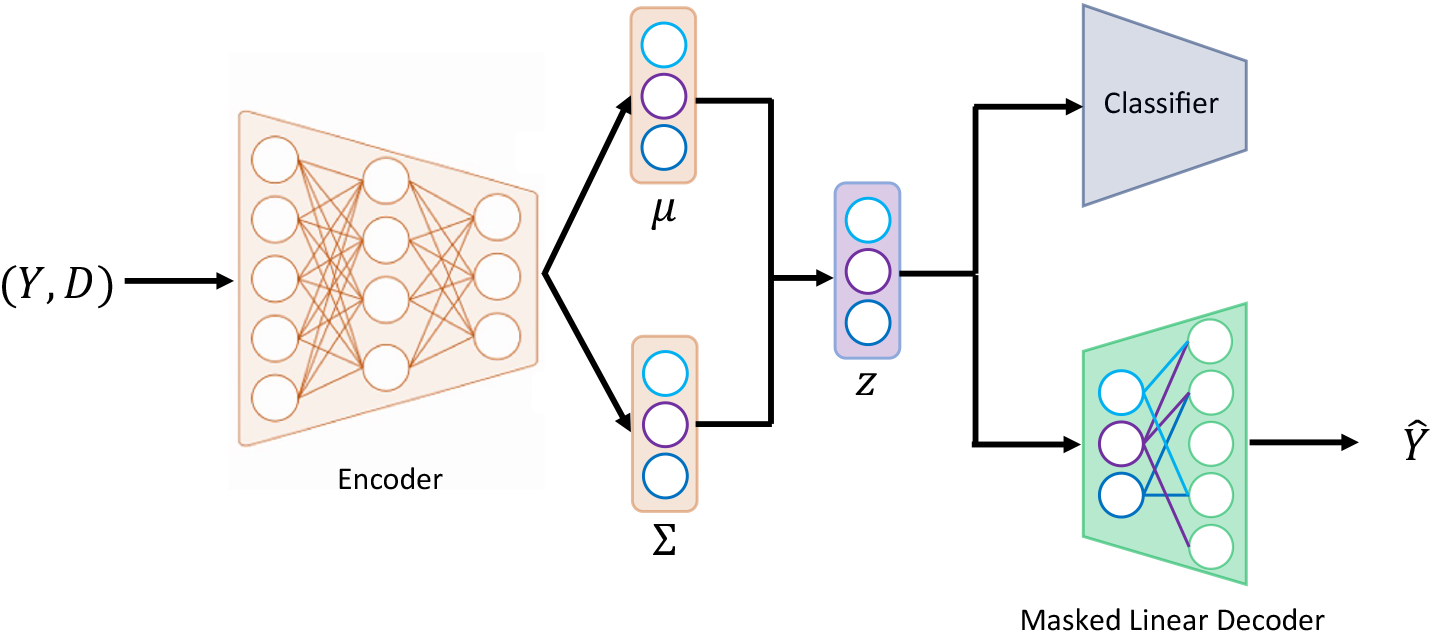
The architecture of EXPORT alternative model in which the ordinal regressor network is replaced with a classifier.

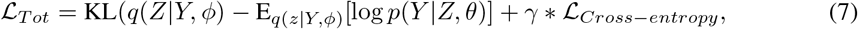

where *γ* is a a coefficient hyperparameter on the cross-entropy loss controlling the effect of classifier in training the overall model. We investigate the latent space learned by this model by analyzing the radiation exposure gene expression data. Similar to original EXPORT model setup, we train the model for 200 epochs and change *γ* as well to see the effect of the cross entropy loss in the inferred latent space. Figure 11 displays the latent space learned by this classification based EXPORT model for different values of *γ*. In Figure 11.D which *γ* = 1*e*6 is used, we can see that the samples are clustered together according to their dosage levels but the ordinality is lost in the latent space. As the ordinality is overlooked in the inferred latent space of this deep learning model and samples with close dosage levels may have very different inferred pathway activities, performing any downstream analysis such as differential pathway analysis lead to inaccurate results. This highlight the need for the ordinal regressor network and cumulative link loss when analyzing the ordinally perturbed transcriptomics data.

**Figure 11:**
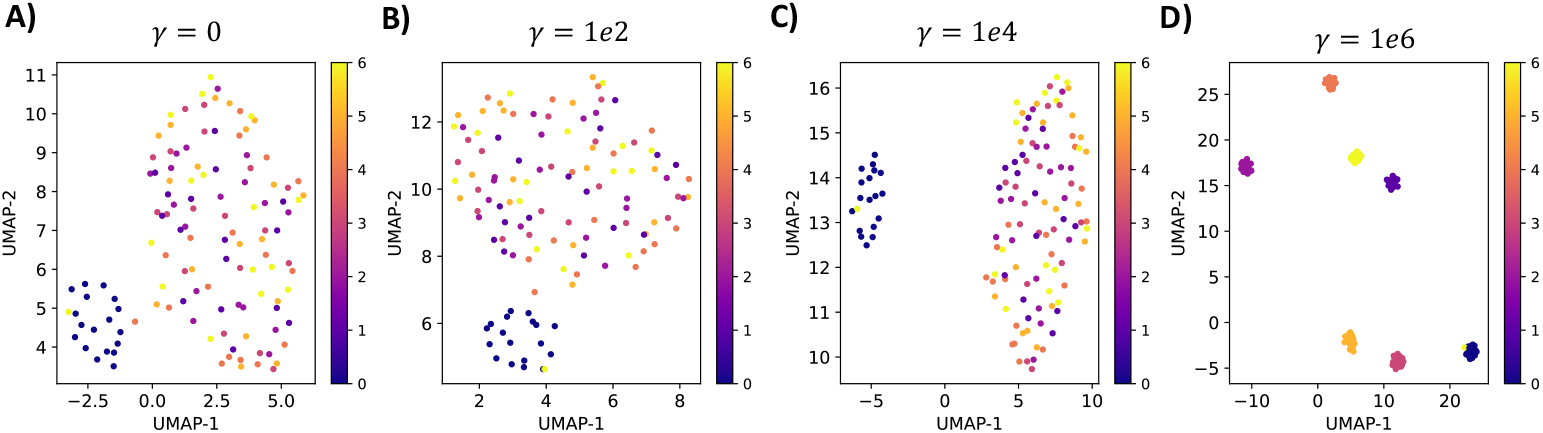
UMAP embedding of latent space learned by EXPORT alternative models with different *γ* hyperparameter values.

## E Training procedure details

Unless specified otherwise in the manuscript methodology section, we used the following hyperparameters for EXPORT: For EXPORT model training we used a learning rate of 1e-4 with the Adam optimizer. The batch size was set to 128 for training EXPORT on TCDD-perturbed snRNA-seq dataset. The KL divergence loss term in EXPORT total loss described in Equation 6 was weighted with a factor *β* = 0.00005. The latent space dimension was defined by the number of biological pathways used for the analysis, plus an additional fully connected node to capture additional variance. Specifically, the following dimensions were used in our experiments:

**Radiation exposure dataset:** 343 KEGG pathways + 1 FC node = 344

**TCDD-perturbed snRNA-seq dataset:** 202 Mouse Wikipathways + 1 FC node = 203

